# An Evidence-based Cognitive Model of Uncertainty during Indoor Multi-level Human Wayfinding

**DOI:** 10.1101/2022.07.27.501728

**Authors:** Qi Yang, Rohit K. Dubey, Saleh Kalantari

## Abstract

Existing computational models lack adequate representation of the uncertainty experienced in human wayfinding tasks. They overly rely on optimized pathing algorithms, which reduces realism and limits insights on human responses to architectural designs. To address this, we developed an empirically grounded model that predicts human wayfinding uncertainty experience. Using data from 28 participants navigating an educational building with varying signage, we constructed the model (Study 1), and validated it with data from 11 other participants (Study 2). We found the wayfinding uncertainty correlated with the time elapsed since seeing the last helpful sign. The cognitive agent based on this model closely replicated human-reported uncertainty levels during wayfinding tasks under different signage conditions. Although the model more closely resembled human behavior compared to a shortest-route algorithm, additional environmental variables and heuristics are needed for better human outcome alignment. Our study showcases that evidence-based cognitive agent modeling can provide nuanced, human-like wayfinding behavior, enhancing the potential for effective computational design evaluation.

## 1. Introduction

Feeling uncertain during navigation is a common everyday experience. Arriving at a complex intersection within an unfamiliar building, we are unsure of which way to go. Sometimes, we are not able to find enough information to know for certain which choice is best; in other cases, we have to choose between conflicting information. Even after deciding and starting a route, we may continue to doubt if we are on the right track. Though such experiences are familiar to most people, the formal definition of wayfinding uncertainty is hard to pin down. Researchers have defined uncertainty in different ways; for example, Anderson and colleagues defined it as a person being “conscious[ly] aware of ignorance” (Anderson et al., 2019). Bar-Anan and colleagues, in contrast, provided a more objective definition of uncertainty as, “lacking information” (Bar-Anan et al., 2009). A more complex perspective was given by Hirsh and Mar, who used the term “psychological entropy” to describe uncertainty as emerging from conflicting perceptual and behavioral affordances, which means there is a gap between what a person is able to perceive about the environment and what the environment actually allows them to do (Hirsh et al., 2012). For the purposes of the current study, we defined uncertainty broadly, as a subjective state of cognitive or affective ambiguity that arises when an individual lacks information, control, or predictability about a particular situation or event. In some cases, uncertainty may be desirable, for example if it enhances creativity and enjoyment, or encourages exploration (Anderson et al., 2019; Bar-Anan et al., 2009; Keller et al., 2020). In most wayfinding contexts, however, it is more likely that uncertainty will lead to problematic navigational outcomes, as well as to negative experiences such as anxiety and frustration (Carleton, 2016; Kalantari, 2016; Martin & Richter, 2015).

A cognitive model simulating realistic human experiences of navigational uncertainty has valuable uses in the fields of environmental psychology, architectural design, and related areas of endeavor. The modeling process allows us to test and validate theories about uncertainty at a more robust and fine-grained level. The resulting validated and predictive model can then serve as a diagnostic tool for helping to identify potential problem-areas in a design before an environment is constructed. Finally, there is an increasing interest in developing “smart” or interactive wayfinding assistive tools (Liu et al., 2022; Zhang et al., 2021). Such tools have to strike the right balance between over-assistance, which tends to annoy users and may lead to over-reliance and a decline in unassisted wayfinding skills, vs. providing too little information. An improved understanding and realistic model of wayfinding uncertainty can help such tools to offer assistance in a timely and precise fashion.

Previous studies of human wayfinding behavior have tended to regard uncertainty as a background condition rather than as a topic of inquiry in itself. This situation is beginning to change in the field, as more researchers become interested in understanding and/or modeling uncertainty. For example, a study by Brunyé and colleagues evaluated how uncertainty in information sources affects human route selection (Brunyé et al., 2015), and work by Keller and colleagues analyzed the types of information sources that people tend to look for when they experience navigational uncertainty (Keller et al., 2020). In 2017, Jonietz and Kiefer presented what might be the first conceptual computational model of wayfinding uncertainty, using a non-deterministic reference system consisting of an agent, an environment, and a route instruction (Jonietz & Kiefer, 2017a). Their work focused primarily on uncertainty in wayfinding situations where instructions that are given to the agent do not fully determine the route. For example, ambiguity can arise from the instruction “turn at the yellow postbox” in an environment where two or more different route choices are available near the yellow postbox. However, this is only one type of possible wayfinding uncertainty, and it does not fully reflect the complex process of searching for signs and other local environmental information to make a navigational decision.

In the current paper, we established a broader conceptual framework for simulating wayfinding uncertainty when the available information (based on directional signs) is conflicting or incomplete. We grounded this interpretation of wayfinding uncertainty on the concept of “oriented search,” which was proposed by Allen and colleagues as a way of understanding situations in which people rely primarily on directional signs for pathfinding through an unfamiliar environment. Such oriented search behavior is typical of wayfinding tasks in large and complex urban facilities (Allen, 1999). We further based the proposed uncertainty formulation on Hirsh and colleagues’ view of competing perceptual and behavioral affordances (Hirsh et al., 2012). Our goal was to include in the model both route-choice uncertainty (when a person is approaching a decision point and has to select a route) and on-route uncertainty (when a person is moving between decision points, for example walking down a long hallway, and is still seeking to confirm that their current route is the correct one). To the best our knowledge, no currently existing wayfinding model has accounted for both of these uncertainty types.

### 1.1. Related Works

Although common wayfinding simulation approaches assume complete spatial knowledge of the agents (Pelechano & Malkawi, 2008), wayfinding simulation is not just a logistics routing problem. It involves subjective behavior under conditions of incomplete information, which often entails human variability and irrationality. Accordingly, in recent years researchers have started to take a greater interest in integrating human perceptive and cognitive processes into the simulation process. One early cognitive agent framework, proposed by Raubal (Raubal, 2001), described the wayfinding process as a continuous interaction between humans and their environment, incorporating feedback loops. This framework implemented an agent capable of making route decisions based on goals, observations of surrounding information, decision rules, and some prior common-sense knowledge. Although this agent assumed perfect observation of signs and utilized simple decision rules (thus ignoring the uncertainty and variability inherent to human wayfinding), it laid the theoretical foundation for realistic wayfinding simulation.

To address the limitations of perfect observation, a subsequent framework introduced ambiguity when mapping wayfinding instructions to route choices (Jonietz & Kiefer, 2017b). Other studies have further diversified decision rules. For example, the Spice cognitive agent framework introduced procedural cognitive processing modules for decision-making (Kielar & Borrmann, 2018). The cogARCH framework incorporated decision heuristics from empirical studies into a behavior tree, enabling the cognitive agent to adopt complex wayfinding strategies. The agent in this framework also had a more realistic vision-based search process, relying on the perception of pre-defined building zones (Gath-Morad et al., 2020). Additional studies have focused on simulating more precise and realistic vision-based human–sign interactions. In these studies, researchers defined a visibility catchment area for the agent, within which the agent acquires information (Becker-Asano et al., 2014; Chen, 2012; Dubey et al., 2021; Maruyama et al., 2017b). This approach allows for a more accurate representation of how humans interact with signs in their visual field.

While these prior works helped to establish the necessary computational groundwork for the cognitive wayfinding agent approach adopted in our current study, most of them did not incorporate empirical measures of uncertainty into the agents’ behavior. One exception to this is Maruyama’s simulation, which identified disorientation areas resulting from a lack of signage (Maruyama et al., 2017a). However, the uncertainty criterion employed in Maruyama’s study was relatively simple, determined by solely the lack of signs, and the validation sample size was limited. Our study aimed to bridge this gap and enhance the realism of indoor wayfinding simulations by using empirical research to identify a broader range of relevant factors that contribute to uncertainty. The resulting model can provide valuable feedback on human responses to assist in early-stage design evaluation.

### 1.3. Research Objectives

We used an evidence-based simulation approach that included three steps. First, we conducted an empirical study to evaluate the impact of several environmental and signage-related features on self-reported uncertainty levels during wayfinding (Study 1). Participants in this initial study were asked to complete wayfinding tasks in a complex building while reporting their uncertainty levels in a continuous fashion. Second, we constructed an uncertainty prediction model based on the high-impact features, and embedded it within a vision-based cognitive agent. Finally, we validated the agent’s uncertainty predictions against unseen human data (that is, data under a different signage condition that was not used in constructing the simulation) (Study 2). In the broadest sense, the objective of this work was to improve our understanding of the relationship between environmental design factors and human uncertainty during navigation, while advancing the field of agent-based navigational modeling via empirical study and validation.

## 2. Study One: Effects of Environmental Elements on Uncertainty during Human Navigation

The first aspect of this project was to conduct a real-world wayfinding experiment to evaluate signage-related environmental features that affected continuous uncertainty experience. We had a fortunate opportunity when our university decided to renovate the directional signs in a large and complex campus building, without changing other environmental features. We collected wayfinding data with different participants both before and after the sign renovations. The post-renovation data was used in Study 1 to evaluate the impact of environmental features on uncertainty experience and to create the predictive model. We then used the pre-renovation data to validate the model under a different signage system (Study 2).

### 2.1. Participants

Thirty-nine participants with normal or corrected-to-normal vision were recruited through convenience sampling (a departmental e-mail list). The inclusion criteria for the study were that the participants should be able to readily complete a series of wayfinding tasks in an indoor environment (including the use of stairs), that they had no significant motor impairments, and that they could readily understand written and spoken English. We collected demographic information on age and gender; these factors were not analyzed as variables in our study, but the data are reported here to help provide a snapshot of the overall participant sample. Data on ethnicity and other demographic variables beyond gender and age were not collected.

Eleven participants conducted the experiment before the sign renovations. Their ages ranged from 17 to 21 (*M* = 19.55, *SD* = 1.29). Among them, two participants were categorized as “Expert” wayfinders (having visited the building a minimum of ten times and expressing confidence in giving wayfinding directions for the building). Nine participants were categorized as “Novice” wayfinders (had previously visited the building no more than once). We had an additional category of “Moderately Experienced” wayfinders for participants who had visited the building between 2–9 times, but none of the participants fell into that category. In this pre-renovation group, 9 participants reported as Female, 2 reported as Male, and 0 reported as Other. These participants comprised the validation group.

Twenty-eight participants conducted the experiment after the sign renovations. Their ages ranged from 18 to 36 (*M* = 21.25, *SD* =3.86) Eight of these participants were categorized as “Expert” wayfinders, 20 as “Novice” wayfinders, and again 0 as “Moderately Experienced” wayfinders. In this post-renovation group, 17 participants reported as Female, 10 reported as Male, and 1 reported as Other. These participants comprised the model-training group.

Prior to the wayfinding tasks, we collected data on each participant’s spatial skills using the Santa Barbara Sense of Direction Scale (SBSOD). These skill levels were evaluated as a variable in our analysis, and were used to introduce agent diversity into the simulation, as discussed in the following sections. We also asked participants to complete a measurement of Spatial Anxiety (Lawton, 1994) and a Mental Rotation task(Vandenberg & Kuse, 1978). Descriptive statistics for these measures across each participant group are presented in Table 1. All participants gave informed written consent prior to the research activities, and the study procedures were reviewed and approved by the Institutional Review Board at **[removed for the purpose of blind review]**.

**Table 1.** Study Participants’ Spatial Skills.

| Study Group | Scale | Minimum | Maximum | Mean | SD |
| --- | --- | --- | --- | --- | --- |
| Pre-Renovation Group | SBSOD | 3.53 | 5.33 | 4.24 | 0.57 |
|  | Spatial Anxiety | 1.38 | 4.50 | 2.83 | 0.99 |
|  | Mental Rotation | 3.00 | 12.00 | 7.73 | 3.38 |
| Post-Renovation Group | SBSOD | 2.40 | 6.20 | 4.09 | 0.98 |
|  | Spatial Anxiety | 1.00 | 4.75 | 2.66 | 0.82 |
|  | Mental Rotation | 0.00 | 13.00 | 7.00 | 3.41 |

*Note: Scale ranges were as follows: SBSOD 1–7; Spatial Anxiety 1–5; Mental Rotation 0–24.*

### 2.2. Measuring Continuous Uncertainty Experience

To measure participants’ self-reported uncertainty levels continuously, we developed a “real-time uncertainty annotation” (RCUA) method, which allowed participants to use pressure on a joystick device to easily indicate greater or lower levels of uncertainty as they completed wayfinding tasks. The validation of this measure can be found in our previous paper on (Yang & Kalantari, 2022), Prior studies have applied various self-reported scales to evaluate uncertainty or anxiety in a broad sense, but these measures require extensive questionnaire responses, and are thus unsuited to continuously measuring uncertainty during task completion (Buhr & Dugas, 2002; Sorrentino, 2008). Another approach is to use indirect measurements of associated objective states such as heart-rate (Keller et al., 2020; Luque-Casado et al., 2013; Zhu et al., 2022); the RCUA method contrasts with such measures in that it is a direct evaluation of subjective experiences of wayfinding uncertainty. This measure allows us to model not only what people *do* in a particular wayfinding context (behavioral reactions or pathing maps), but also what their subjective *experiences* are likely to be when they encounter certain environmental features.

The hardware used in this instrument included a Nintendo Wii Nunchuk Controller paired with an 8BitDo Wireless USB Adapter. The Nunchuk controller was selected because it has a large analog stick and is convenient for users to hold and push while standing. We developed a script in Python that documents the real-time input of the controller in terms of the amount of force exerted in any direction from center. Prior to the navigational tasks, participants were instructed to increase their pressure on the joystick when they were feeling greater navigational uncertainty, and they were provided with descriptive examples of what kind of experiences would entail “no uncertainty” (completely certain of the path) and “highest uncertainty” (completely confused about the path). The real-time force exerted on the joystick was recorded by the Python script on a scale of 0 (no force) to 1 (maximum force).

During the real-world wayfinding tasks, participants carried the RCUA joystick and used it to report uncertainty levels while a researcher followed at approximately 5–10 ft. distance. The researcher carried a Tablet (Microsoft Surface Pro) that recorded the wireless data from the joystick and simultaneously ran the **[removed for the purpose of blind review]** App previously developed by our team. The researchers used this app to manually track the participants’ trajectories through the building using timestamps on a floorplan. The participants and the following researcher also wore cameras (GoPro Max), positioned at chest level, to document the wayfinding process (Figure 1).

**Figure 1.**
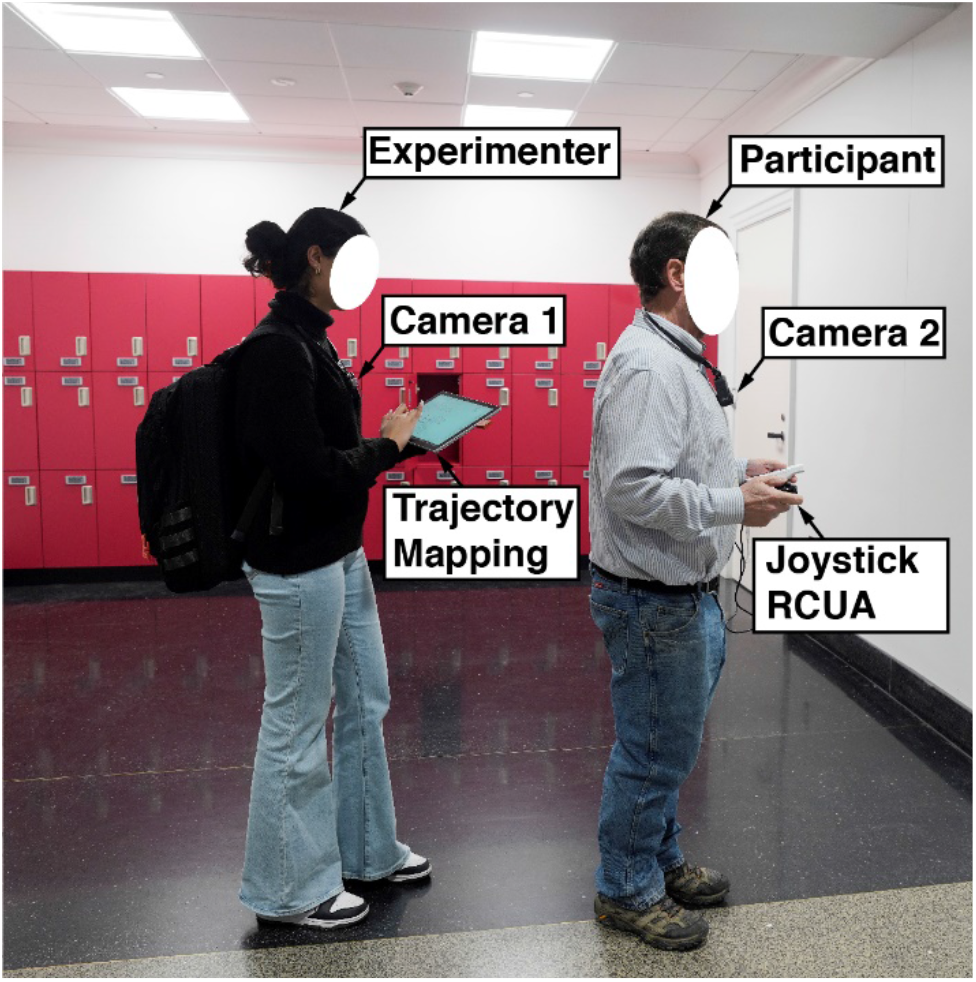
Experiment Setting for Real Time Uncertainty Annotation (RCUA) during Wayfinding Tasks (During Data-collection the Researcher Trailed the Participant by at Least 5 Feet)

### 2.3. Procedure

Sessions were conducted for one participant at a time. Prior to the experiment sessions, participants were asked to complete an online survey that collected the demographic information and the spatial skills measurements. When participants arrived in the lab for their sessions, they were first trained to use the joystick for reporting uncertainty experiences, and were then asked to complete the wayfinding tasks. This involved finding seven specific destinations within the building in a set sequence. The next destination was given only when the previous one had been found. At the start of each individual wayfinding task, the participants were reminded to use the joystick to continuously report their uncertainty level.

### 2.4. Wayfinding Task Design

The experiment was conducted in the **[deleted for the purpose of blind review]** building, which is large and complex and has a reputation on campus for wayfinding difficulties. We designed the seven wayfinding tasks to include a variety of environmental features. Three of the tasks involved moving between different building levels (floors), while the other four tasks could be completed by traveling on the same level. The tasks had varying degrees of difficulty, with different lengths and sign coverage. All of the tasks included navigating at least two intersections, which were regarded as likely uncertainty-inducing points (Hirsh et al., 2012). The seven tasks formed a loop (Table 2 and Figure 2), and the starting task was randomized for each participant to eliminate learning effects in the aggregate data. To minimize confounding variables, participants were not allowed to go outside of the building or ask other people for help. We also stipulated that participants could not use the building’s elevators, since the variable elevator traffic would affect the task completion time.

**Table 2.** Description of the Wayfinding Tasks Used in the Study.

| **Task Number** | **Start Location** | **End Location** | **Type of Wayfinding** | **Shortest Path (in Meters)** |
| --- | --- | --- | --- | --- |
| 1 | MVR Cafe | Room T222 | Single-level | 38.45 |
| 2 | Room T222 | Room T320C | Single-level | 41.32 |
| 3 | Room T320C | Room 1250 | Multi-level | 51.76 |
| 4 | Room 1250 | Room 1106 | Single-level | 103.89 |
| 5 | Room 1106 | Room G151 | Multi-level | 58.52 |
| 6 | Room G151 | Room HEB T70 | Multi-level | 82.43 |
| 7 | Room HEB T70 | MVR Cafe | Single-level | 75.62 |

**Figure 2.**
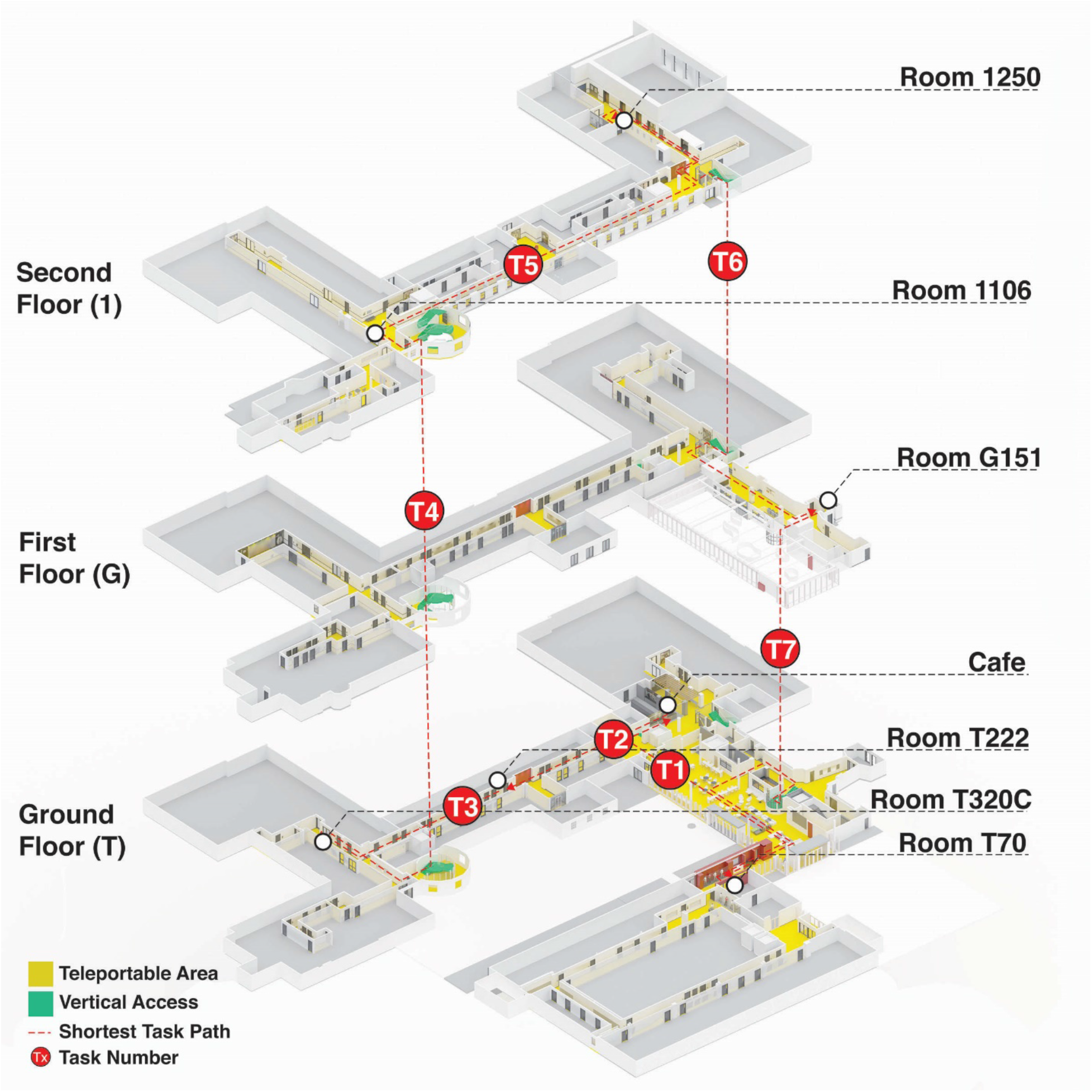
Seven Wayfinding Tasks in Three Floors

### 2.5. Modeling the Environment

We digitized the physical environment to extract and analyze features related to uncertainty experiences. First, we created a digital model of the building layout using the software tool Rhinoceros 6.0 (McNeel & Others, n.d.). Next, we surveyed signs in the building that provided directional information to pedestrians. We collected the information from these signs and organized it into a structured format called JSON files, which allowed us to analyze the types and locations of the signs in the digital model. To better understand the impact of route alternatives on uncertainty, we divided the walkable areas of the building into distinct zones. The areas in which participants had to make key route choices were categorized as “decision-making” zones; and the remaining areas, including long corridors, staircases, and dead ends, were classified as “non-decision” zones. The total number of available route choices in each zone was identified and associated with the zone’s index (Supplementary Material, Figure S1).

As part of identifying relevant environmental features from the human participants’ wayfinding data, we created an agent in the digital model that precisely followed each participant’s trajectory and simulated the participant’s visual perception. This trajectory was sampled at 5 Hz. We defined the agent’s field of view as 120 degrees, and used 10m as the isovist radius (the maximum distance the agent could visually perceive information on a sign). The choice of a 10m visual radius was based on reasonable average maximum viewing distance relative to the size of the physical signs in our test building, both pre-renovation and post-renovation.

### 2.6. Choice of Environmental Features

We approached the issue of features relevant to wayfinding primarily from the work of Dubey (Dubey et al., 2021), who influentially categorized such information sources into five groups: signage, space, memory, crowd, and history. The “signage” category describes information gained from all types of human-made guidance materials, including wall-signs, maps, or textual directions such as “turn left at the fountain.” The “space” category includes environmental elements such as geometric form, lighting, obstacles, landmarks, and available pathways. “Memory” includes previous spatial knowledge of the environment or similar environments; “crowd” describes the influence of nearby people’s behavior on wayfinding choices; and “history” serves as a catch-all term for individual preferences such as the propensity for risk-taking when selecting unknown routes. Much of the recent literature in wayfinding research has adopted this framework—for example by seeking to identify how people prioritize these different categories of information in various environmental contexts, or by evaluating the different cognitive strategies associated with each category of navigational information (Hölscher et al., 2006; Kim, 1999; Rounds et al., 2020; Sharma et al., 2017)

Under the signage category, we evaluated three relevant features. The “Number of Visible Signs” was measured by adding up the total number of signs within the isovist area (i.e., signs that the participant could read at any given time). The “Number of Helpful Signs” was the subset of visible signs that contained directional information related to the current wayfinding task. “Elapsed Time Since Helpful Sign” measured how much time had passed since the last helpful sign was within the agent’s isovist area. For the spatial category, we extracted 16 metrics within the isovist representational framework, which include important aspects of how humans experience their surroundings, such as the shape of the visible area and its skewness compared to the direction of motion (Arabacioglu, 2019; Wiener & Franz, 2015). These metrics were extracted using the DecodingSpaces toolbox (Abdulmawla et al., n.d.), and are presented in more detail in the supplementary material (Table S1). We also included the “Number of Route Choices,” measured by the number of available corridors connected to the current wayfinding decision zone.

Under the memory and history categories, previous studies have shown that participants’ previous spatial knowledge of the environment, as well as their self-reported sense of direction as measured by the SBSOD, can affect wayfinding performance (Kuliga et al., 2019). These variables were operationalized by our division of participants into “experts” vs. “novices” in relation to their prior knowledge of the building, and by the SBSOD scores. We did not include any variables related to Dubey’s “crowd behavior” category, since this would be quite difficult to measure empirically and to add to the cognitive agent simulation (future work on crowd variables is highly recommended, but would likely entail its own extensive study). We also omitted personality factors beyond the SBSOD that might be related to decision heuristics, such as risk-taking proclivity. The full list of factors evaluated in the current study is summarized in Table 3.

**Table 3.**
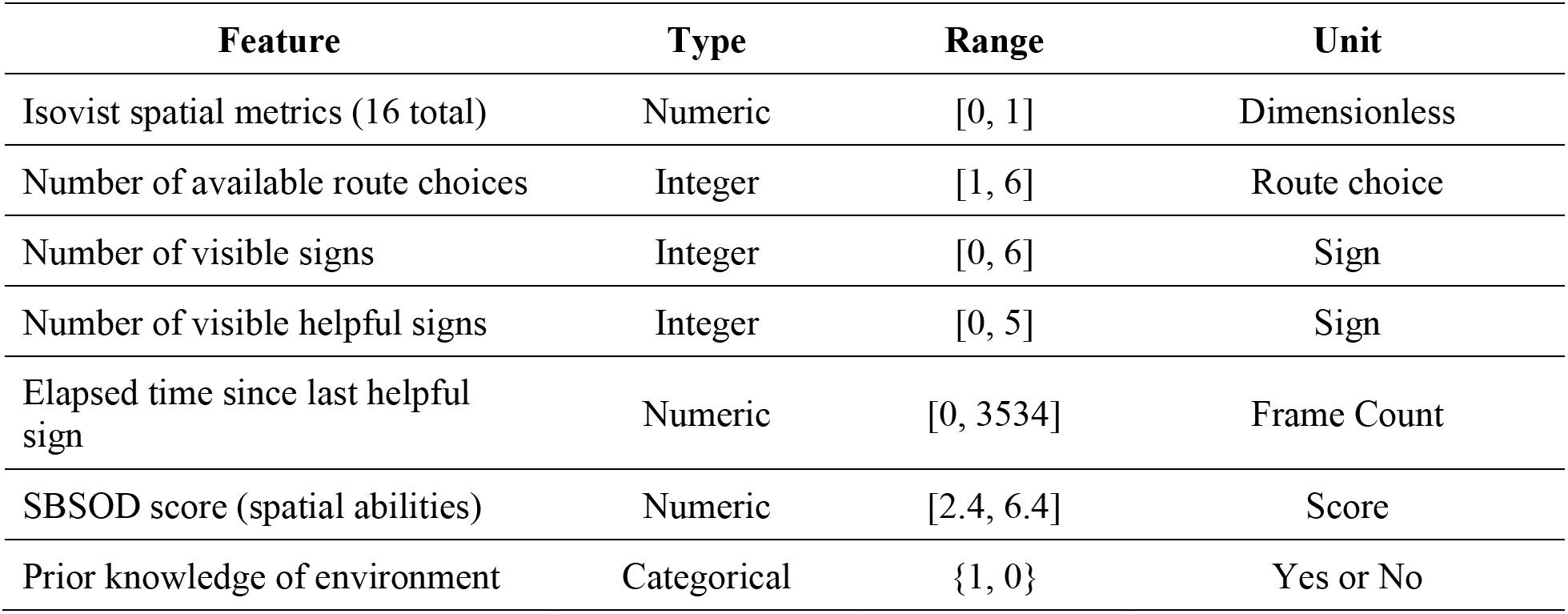
Identified Features that Affected Wayfinding Uncertainty.

| **Feature** | **Type** | **Range** | **Unit** |
| --- | --- | --- | --- |
| Isovist spatial metrics (16 total) | Numeric | [0, 1] | Dimensionless |
| Number of available route choices | Integer | [1, 6] | Route choice |
| Number of visible signs | Integer | [0, 6] | Sign |
| Number of visible helpful signs | Integer | [0, 5] | Sign |
| Elapsed time since last helpful sign | Numeric | [0, 3534] | Frame Count |
| SBSOD score (spatial abilities) | Numeric | [2.4, 6.4] | Score |
| Prior knowledge of environment | Categorical | {1, 0} | Yes or No |

### 2.7. Statistical Methods

Previous studies have shown associations between our evaluated factors and wayfinding performance, but we found that there was not much literature analyzing the relationships between these features and *uncertainty*. In the absence of robust prior findings, our statistical approach assumed potential linear relationships and weak interaction effects. We therefore used linear correlation tests to evaluate the strength of association between reported uncertainty levels and each studied feature, and selected the features with stronger correlations to fit a multiple linear regression model. For the Prior Knowledge of the Environment variable, we partitioned the dataset into “experts” and “novices” and used a one-way ANOVA against the participants’ average uncertainty levels.

### 2.8. Results of Study 1

The isovist spatial metrics had little impact on participants’ uncertainty. Most of these metrics had a correlation of |r| < 0.02, which is considered negligible. The two variables of Circularity and Elongation of the isovist space had a slightly stronger positive correlation of r = 0.06, but this is still considered close-to-trivial. The overall Number of Visible Signs also had a negligible correlation with uncertainty at r= -0.03. We found a very weak positive correlation between the Number of Route Choices and uncertainty (r = 0.08) and a very weak negative correlation between the Number of Visible Helpful Signs and uncertainty (r = -0.1). There was a greater correlation between the Elapsed Time Since Helpful Sign and uncertainty (r = 0.21).

As for the individual factors, SBSOD scores were found to have a moderately strong negative correlation with uncertainty (r = -0.45), as would be expected for participants with greater spatial skills. The ANOVA test for Prior Knowledge of the Environment also showed significant association of experience with reduced uncertainty levels (F(1, 26) = 5.731, p = 0.024). These results are summarized in Table 4.

**Table 4.** Correlation Coefficients between Perceived Uncertainty and Variables.

| Features | All | Expert | Novice |
| --- | --- | --- | --- |
| Number of visible signs | -0.03* | 0.03* | -0.03* |
| Number of visible helpful signs | -0.1* | -0.22* | -0.13* |
| Number of route choices | 0.08* | 0.18* | 0.04* |
| Elapsed time since helpful sign | 0.21* | 0.72* | 0.19* |
| SBSOD | -0.45* | -0.16 | -0.42 |

*Note: alpha = 0.05; \* significant*

On the basis of these findings, we excluded all of the isovist metrics as well as the Number of Visible Signs from our multiple linear regression model. (The Number of Visible Signs was excluded both due to its negligible correlation and to its potentially complex interaction with Number of Helpful Signs. An overall large number of signs might reduce uncertainty if the signs are helpful, but if there many unhelpful signs then they could plausibly have an effect of increasing uncertainty. This may help to account for the negligible correlation in the findings.) The multiple linear regression model therefore used Number of Visible Helpful Signs, Number of Route Choices, Elapsed Time Since Helpful Sign”, SBSOD Score, and Prior Knowledge of Environment as fixed effects and participant number as random effects. The marginal r-squared for the model was 0.196, and the conditional r-squared was 0.388. The parameters of the model are shown in Table 5. This model was used to develop the agent-based simulation of wayfinding uncertainty in the remainder of the project.

**Table 5.** Parameters of the Multiple Linear Regression Model.

|  | Coefficient | Standard Error | *t* | *p* |
| --- | --- | --- | --- | --- |
| Intercept | $6.046 \times 10^{-1}$ | $1.575 \times 10^{-1}$ | 3.838 | 0.0007 |
| Number of visible helpful signs | $3.063 \times 10^{-1}$ | $5.723 \times 10^{-4}$ | 53.522 | <0.0001 |
| Number of route choices | $-4.902 \times 10^{-1}$ | $2.090 \times 10^{-3}$ | -23.459 | 0.0001 |
| Elapsed time since helpful sign | $2.436 \times 10^{-1}$ | $2.457 \times 10^{-6}$ | 99.150 | <0.0001 |
| SBSOD | $-1.030 \times 10^{-1}$ | $3.888 \times 10^{-2}$ | -2.646 | 0.0138 |
| Prior knowledge of environment | $-1.462 \times 10^{-1}$ | $8.270 \times 10^{-2}$ | -1.768 | 0.0892 |

## 3. Cognitive Agent Development

In this section, we present in detail the computational model of the cognitive agent grounded in the above experimental findings. The goal of this agent is to produce human-like uncertainty during wayfinding in an unfamiliar environment. We begin by discussing the agent’s visual perception model, along with the signage and environmental model used in the proposed computational framework. Both are crucial for reproducing the results and performing realistic validation. We then discuss the formulation of an agent that incorporates both wayfinding heuristics and uncertainty modeling during navigational tasks. The Unity3D game engine (https://unity3d.com/) was employed to develop the environmental model and simulate the agent’s movement behavior.

### 3.1. Visual Perception Model

Visual cues in the environment are the primary source of information for most people as they seek to complete wayfinding tasks (Schinazi et al., 2016). (Our work did not consider the specific wayfinding needs of the visually impaired, which is a crucial topic but is beyond the scope of the current research.) To realistically simulate visual-based interactions between agents and their environment, a human-like visual perception model is needed, with an emphasis on evaluating the visibility of directional signs and considering dynamic occlusions of such signs (Moussaïd et al., 2011). We developed a basic perceptual-based agent–signage interaction system for this purpose. Our approach in this area was inspired by Dubey’s similar prior work (Dubey et al., 2021). To account for typical human neck rotation when searching for nearby environmental cues, our model defined the agents’ horizontal field of view as 120 degrees (Fisher et al., 1987). The full interior height, floor to ceiling, was assumed to be visible within this field of view. The distance from which signs can be read (isovist radius) was set to 10 meters. These parameters can be adjusted by the model’s users as needed, for example in environments with very large or very small signs.

### 3.2. Signage and Environment Model

The basic environmental model consisted of a 3D building floorplan imported to and augmented (adding stairs between floors) in the Unity 3D engine. Individual directional signs were identified in the simulation by their attributes and a list of relevant goal locations (Figure 3). A sign was regarded as visible to an agent when it was within that agent’s field of view and there was no occlusion, as determined by a dynamic visibility check (Dubey et al., 2021). The cognitive agent’s wayfinding behavior was implemented using a navigation graph. Each intersection (i.e., decision point) acted as a node in the navigation graph, with paths (corridors or hallways) connecting two nodes. Decision points were identified computationally based on the method suggested in previous study with human in the loop (Dubey et al., 2020).

**Figure 3.**
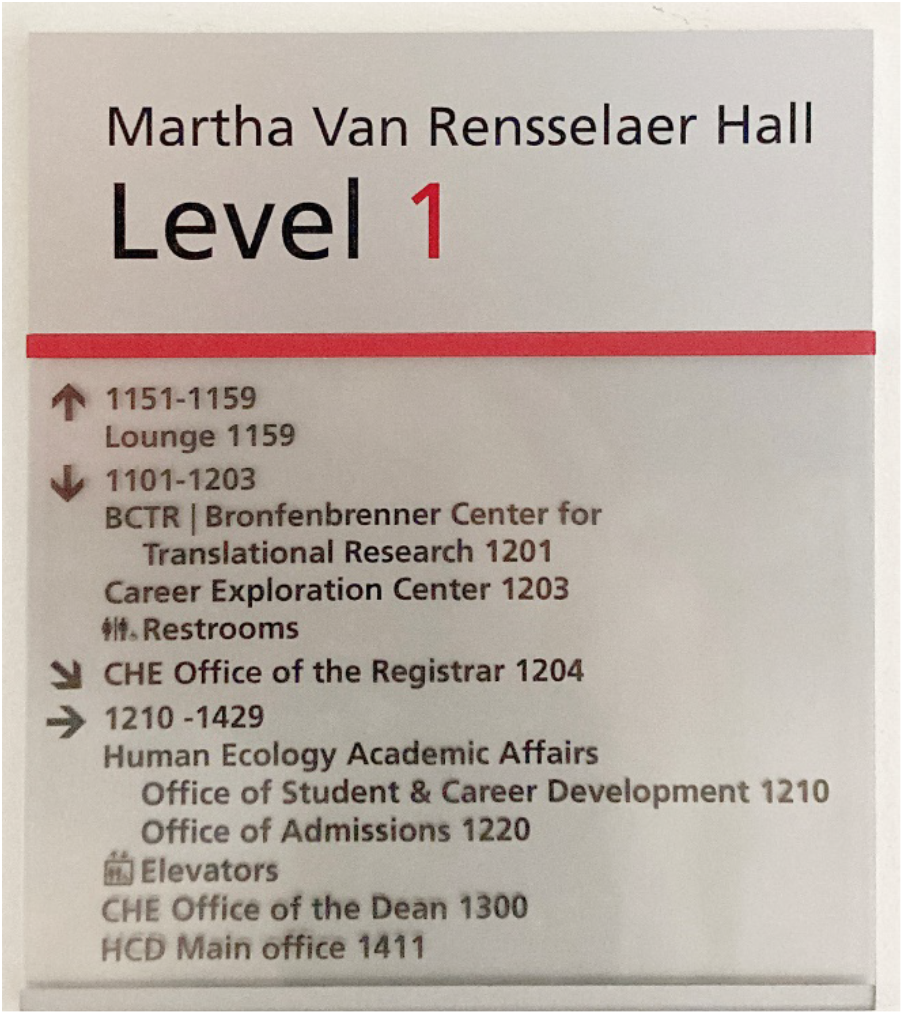
One Example of the Directional Signs in the Building

### 3.3. Simulating Natural Movement

The model’s goal was to simulate a human’s natural wayfinding movement pattern without a pre-determined or optimized path (that is, a visitor’s exploration behavior while navigating through an indoor environment with imperfect information). We used a partial isovist approach—that is, a constrained visible area limited by the 120-degree field of view and the surrounding walls and objects—and identified a “drift” point in the centroid of the agent’s field of view toward which the agent moves by default. The centroid is a mathematical center point of the visible space, as determined by the contours of the visible area (Figure 4). The process of identifying the isovist space and centroid is continuous as the agent movement changes the drift position dynamically. As uncertainty is reduced, the amount of drift is reduced and thus the agent moves more directly toward its accurate destination. Under conditions in which the agent is entirely certain about its path, it follows the shortest-path-based locomotion toward the destination.

**Figure 4.**
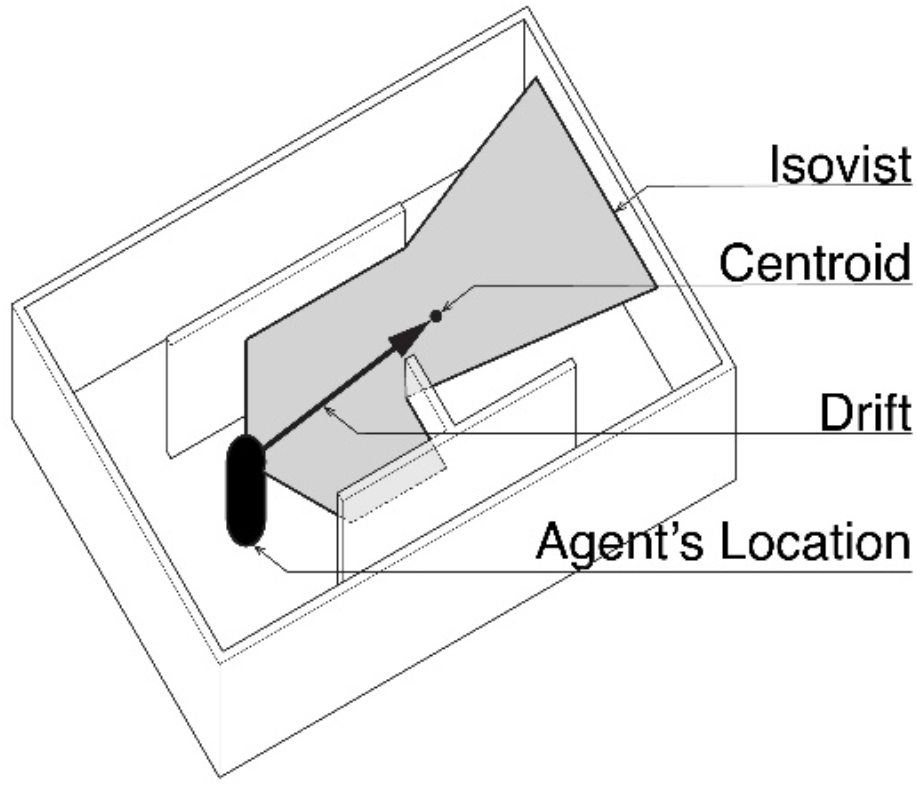
Visualization of An Agent’s Natural Movement for Three Time-Steps Using Isovist Drift Measures.

In addition to drift induced by uncertainty, we also incorporated wayfinding heuristics into the simulation model. Tenbrink and colleagues developed a comprehensive list of such heuristics; that is, strategies used by people when attempting to complete a navigational task (Tenbrink & Wiener, 2007). In our model some of these well-established heuristics were used as meta-strategies guiding the overall direction of drift/search behavior. These heuristics included:

- **Floor Strategy (Strategy 1)**: Finding a route to the destination floor was prioritized irrespective of the horizontal position of the goal (i.e., the agent seeks to find a way to reach the correct floor before searching for the specific destination). Agents are assumed to know the destination floor level, presumably as information included in the destination label itself or as part of the room number. Signs indicating access to the destination floor are prioritized over all other environmental inputs.
- **Following continuously marked room numbers (Strategy 2)**: If the environment includes signs that present ordered room numbers (for example, room 312, 313, 314), then the agent will prioritize this information and move in the direction that is numerically closest to the destination room number.
- **When in doubt follow your nose (Strategy 3)**: In an unfamiliar environment, when people are in doubt, it has been observed that they tend to follow as straight a route as possible with minimal angular deviations (Dalton, 2003; Meilinger et al., 2014). Agents will decide on a path and then generally tend to move in the direction that they have previously started, except as impacted by the other heuristics and uncertainty-induced “drift” behavior. (In this context, following prior usage in the wayfinding literature, “follow your nose” is interpreted to mean moving straight ahead).

In terms of how these heuristics are applied, Equation 1 was used to calculate the vertical distance (number of floors) between the origin and destination floor. The first letter of room numbers on the detected directional signage is compared with the first letter of the destination room number assigned to the agent at the beginning of a wayfinding task. (The heuristic assumes that, following common convention, all rooms on the same floor begin with the same number or letter.) If the room is on a different floor from the current one, then the agent enters into search behavior to find a floor transition (escalator/lift/stairs). If the room is on the same floor, then the agent begins searching the current floor for the destination.

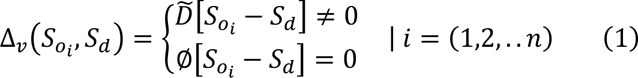

The function Δ_!_(Vertical distance) outputs the state-decisions made by Strategy S1. *S*_*oi*_ is a list of room numbers observed in the agent’s exploration of the current floor. *n* is the number of signs observed. *S*_*d*_ is the destination assigned to the agent. If the first letter/number of the destination room does not match one of the visually observed room numbers, then the agent is assigned a new temporary sub-goal of *D* to search for a transition point (e.g., lift, stairs, or escalator); otherwise, it continues with the same-floor search (designated Φ) with an assumption that the destination is on the same floor.

Strategy *S2* comes into play once the agent believes it is on the correct floor. This strategy is implemented by evaluating the horizontal distance between the current position and the destination when a series of visible room numbers is identified on consecutive signs. Equation 2 is used for this purpose. The agent turns and moves toward the direction that minimizes its distance to the destination based on the ordered-room-number hypothesis.

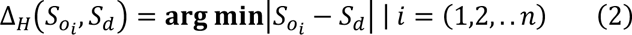

The function Δ_*H*_ (horizontal distance) outputs the overriding directional decision made by *S2*. A simple subtraction operation is performed between the currently observed room number in the series and the destination room number. If the next observed room number results in a greater difference, then the agent turns around and moves in the opposite direction.

Finally, heuristic Strategy *S3* is present in the model as an implicit parameter. In the absence of any new input from observed numerical signs that would merit a floor-change or route-change, the agent will continue to travel in its previously selected direction, apart from the impact of uncertainty-induced drift and additional uncertainty mechanics as described below.

### 3.4. Wayfinding Uncertainty Model

Uncertainty levels during navigation are continuous and vary throughout the course of a wayfinding task. We use the model derived from the empirical findings of Study 1 to continually predict an agent’s uncertainty experience. The model identifies five parameters that influence uncertainty during indoor wayfinding, which we have outlined as follows: *X*_1_ is the number of route choices at a decision point (corresponding to *U_r_*), *X*_2_ is the number of visible helpful signs, *X*_3_ is the elapsed time from seeing the last helpful sign (corresponding to *U*_a_), *X*_4_ is the agent’s score on the Santa Barbara Sense of Direction Scale (in the simulation, we assigned the agents the same SBSOD scores as the humans in the validation set.), and *X*_5_ is the prior spatial knowledge of the participant (“Expert” or “Novice” in relation to the current environment, which could also be used to create agent variability). Based on this empirical model (Table 5), the overall uncertainty of an agent ∇ during wayfinding is shown in Equation 3. Here, *β*_0_ is the intercept from the regression model, and *β*_1_ through *β*_5_ are the empirically determined regression coefficients for each environmental variable.

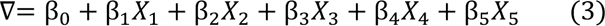

*X*_1_ is assigned to zero when the agent is outside a decision-making zone, under the assumption that route-choice uncertainty does not arise in those portions of the building.

### 3.5. Impact of Uncertainty in the Cognitive Agent Wayfinding Model

Figure 5 presents the overall structure of the proposed agent-based model integrating the effect of uncertainty states. We modeled the agents’ behavior as a finite state machine, meaning that each agent can be in exactly one of a finite number of states at any given time. The agent can change from one state to another in response to environmental input. The proposed model includes five agent states, which are mediated by a combination of the high-level heuristics and the agent’s current uncertainty level:

- *Explore:* In this state, the agent explores the environment, exhibiting a moderately high degree of drift, seeking for directional cues and using its natural movement as discussed in Section 3.3.
- *Sub-goal:* In this state, the agent is assigned a temporary sub-goal of finding a floor transition (staircase/escalator/lift) and performing a shift in floor level. This state occurs when the agent has determined that the destination is not on the current floor.
- *Decision-node*: In this state, the agent is inside a decision zone and must choose between branching available routes. The choice of route depends on information available in the environment. If there is a helpful sign in visual range, then the agent selects the directional information perceived from that helpful sign. In the absence of any helpful sign, the agent selects the route with the longest visual sight range (i.e., line of sight). If multiple unknown routes have the same sight range, then the agent will select randomly between the routes which were not taken previously. (The agent maintains a short-term memory using a navigation graph of routes taken during the wayfinding task.)
- *Execute:* In this state, agent navigates directly toward the identified destination.

**Figure 5.**
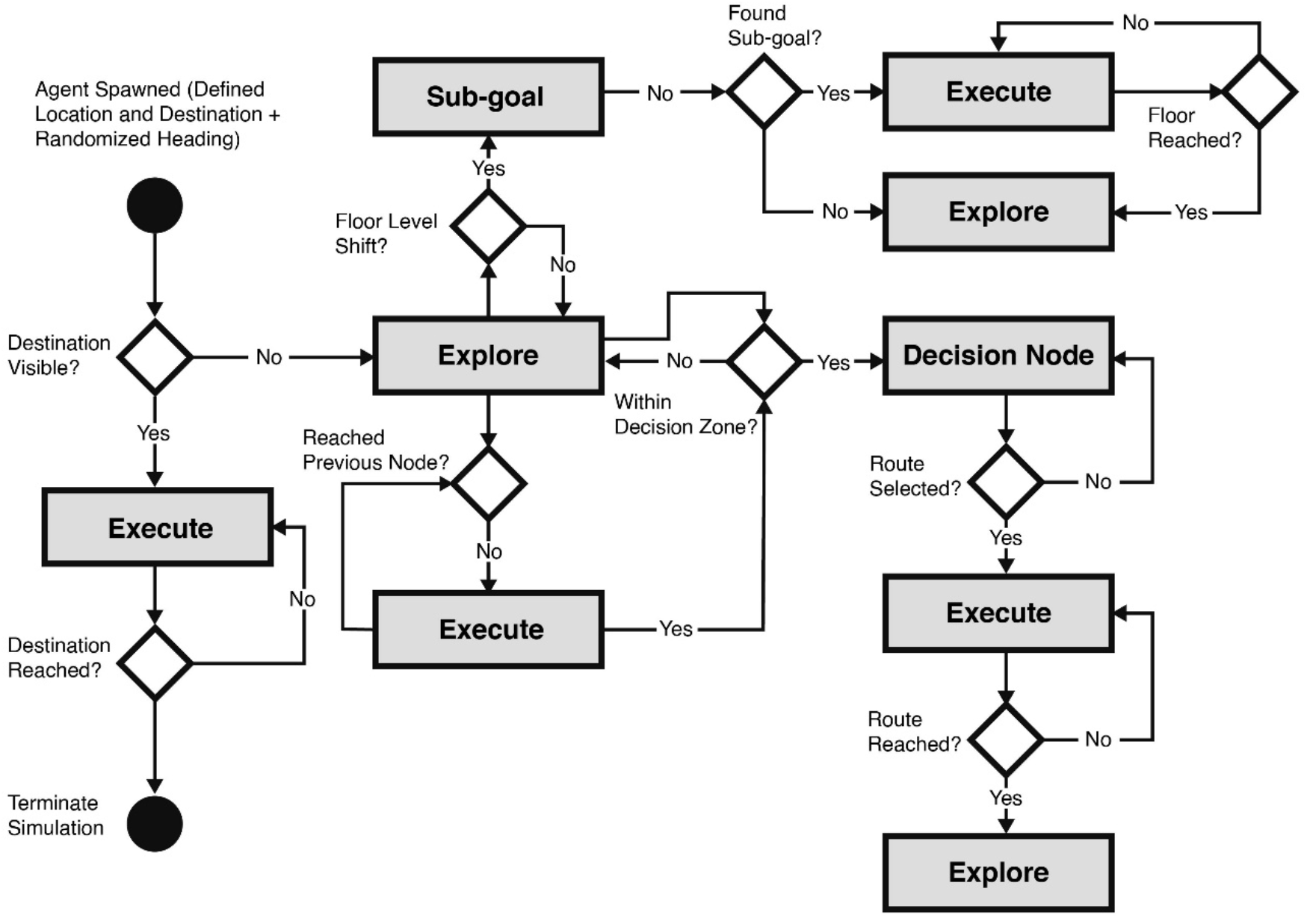
State Machine Diagram Depicting Agents’ Wayfinding Behavior

## 4. Study Two: Validating the Cognitive Model of Uncertainty during Wayfinding

To validate the simulation model we compared its predictions against data from human participants that were not used in the model-creation process. The validation data was collected in the same building and using the same wayfinding tasks, but under a different signage design and with a different group of participants. As discussed in Section 2, there were 11 participants in the validation study. The data-collection process for these participants was identical to that used in the collection of the model-training data. For the purpose of outcome comparisons, we simulated 11 cognitive agents and assigned to them the same starting positions and wayfinding tasks as the human participants. In addition to our proposed uncertainty-based agent, we repeated this process with two baseline computational models: a simple heuristic agent and random agent. The heuristic agent used a simplified uncertainty model based only on the number of route options at a decision point (normalized between 0 to 1, with a greater number of route choices producing greater uncertainty). For the random agent, uncertainty levels at each computation point were assigned by generating a random value between 0 and 1.

To better visualize the wayfinding outcomes, we created path heatmaps showing the uncertainty levels averaged across all 11 participants/agents in a particular category (that is, human agents vs. cognitive agents vs. heuristic agents vs. random agents). To create these heatmaps, the navigation areas were divided into 1 square-meter cells. Each cell value was determined by averaging the uncertainty values measured on participants’ or agents’ trajectories within a 3m radius of the cell. The aggregated heatmaps across all wayfinding tasks are shown in Figure 6, and disaggregated heatmaps for each individual task are presented in the supplementary material’s Figure S2.

**Figure 6.**
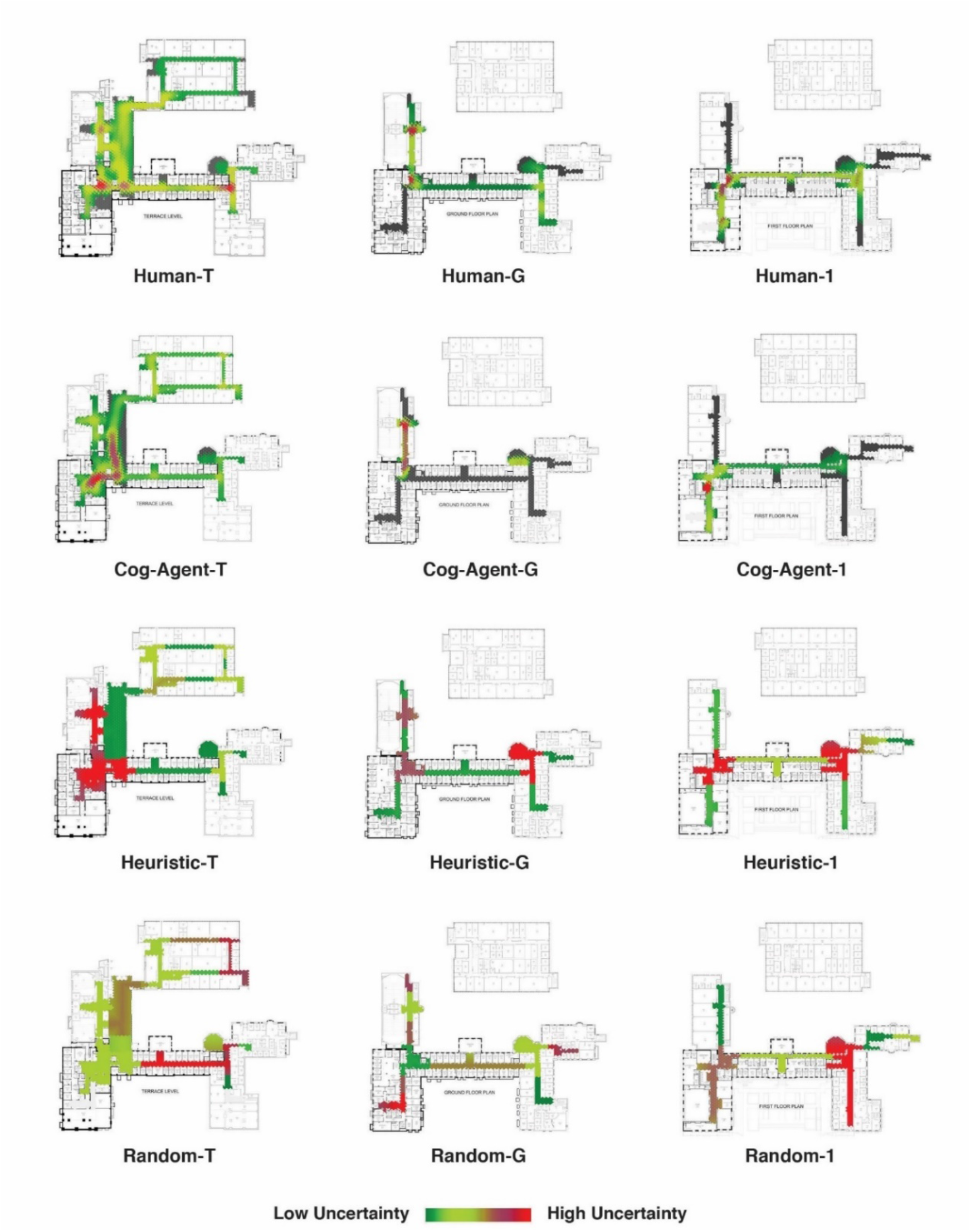
Uncertainty Experienced by Human Participants, the Proposed Uncertainty-based Cognitive Agent, a Simple Heuristic Agent, and a Random Agent, Across All Wayfinding Tasks (T, G, and 1 Refer to the Building Floors) Note: Grey areas indicate potential paths that were not traversed by human and the agent.

In addition to evaluating the distribution of uncertainty levels across the building’s space via the heatmaps, we also used the average path distance as a metric for comparing wayfinding performance. In this metric, we made a three-way comparison of the wayfinding performance of the human participants vs. our uncertainty-based agent vs. the A* shortest-path heuristic agent (Dijkstra, 1959). The average distance traveled by agents in reaching a destination is a commonly used metric in spatial wayfinding literature (Gath-Morad et al., 2022).

### 4.1. Results for Uncertainty Similarity

Comparing uncertainty similarity for two different wayfinding trajectories is a challenging task. The uncertainty depends on multiple factors, such as the number of visible signs, which may vary even with slight route alterations, and different agents may remain within the same spatial region for different amounts of time. For this reason, we took the approach of evaluating the spatial distribution of uncertainty levels, averaged across all participants or agents, as visualized in our uncertainty heatmaps. To make a quantitative comparison between these heatmaps, we used the Dynamic Time Warping (DTW) algorithm, which measures the similarity between two temporal sequences that may or may not have equal lengths (Vintsyuk, 1968). The results indicated that the uncertainty levels reported by the human participants were closely similar to those predicted by our cognitive agent, and that this similarity was not found for the simple heuristic agent or for the random agent (Figure 7). This indicates that our proposed computational model was able to successfully capture the environmental factors that influenced human uncertainty under a novel signage regime.

**Figure 7.**
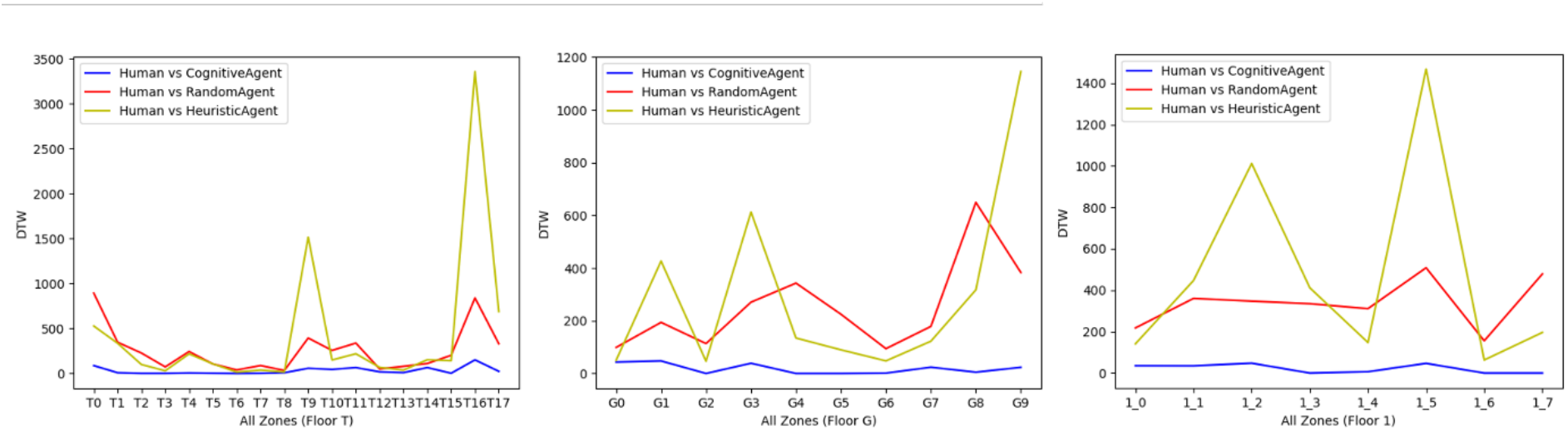
Dynamic Time Warping Comparison of Human Wayfinding Uncertainty vs. the Proposed Uncertainty-based Cognitive Agent, a Simple Heuristic Agent, and a Random Agent, for Each of the Three Floor Levels (Higher DTW Scores Indicate Reduced Similarity)

### 4.2. Results for Average Path Distance

Table 6 and Figure 8 show our findings regarding the average route distances per wayfinding task for the human participants, the proposed uncertainty-based agents, and the A* shortest-path agent. For the human participants and the uncertainty-based cognitive agents, we averaged 11 iterations with different sequences of tasks, as described above. The A* agent outputs a single shortest distance for each task.) The results of this comparison varied greatly among the different wayfinding tasks. We found that the travel distance of the uncertainty-based cognitive agents were most similar to the humans in wayfinding tasks 3, 5, and 6. In these tasks the A* agent’s travel distance was also similar to the humans and the uncertainty-based agent, albeit slightly shorter. In tasks 1, 2, and 7, our uncertainty-based agent took a much longer path than either the humans or the A* agent. In task 4, the humans took a much longer path, while the uncertainty-based agent and the A* agent took shorter paths.

**Table 6.** Average Travelled Distance of Human, Cognitive Agent, and A* Agent.

| Task ID | Cognitive Agent | Human Participants | A*Agent |
| --- | --- | --- | --- |
| 1 | 454.75 | 116.78 | 95.23 |
| 2 | 666.53 | 121.02 | 44.12 |
| 3 | 135.52 | 84.03 | 37.64 |
| 4 | 61.16 | 271.55 | 54.18 |
| 5 | 154.01 | 136.71 | 94.41 |
| 6 | 105.01 | 145.14 | 70.34 |
| 7 | 655.05 | 195.06 | 72.16 |

**Figure 8.**
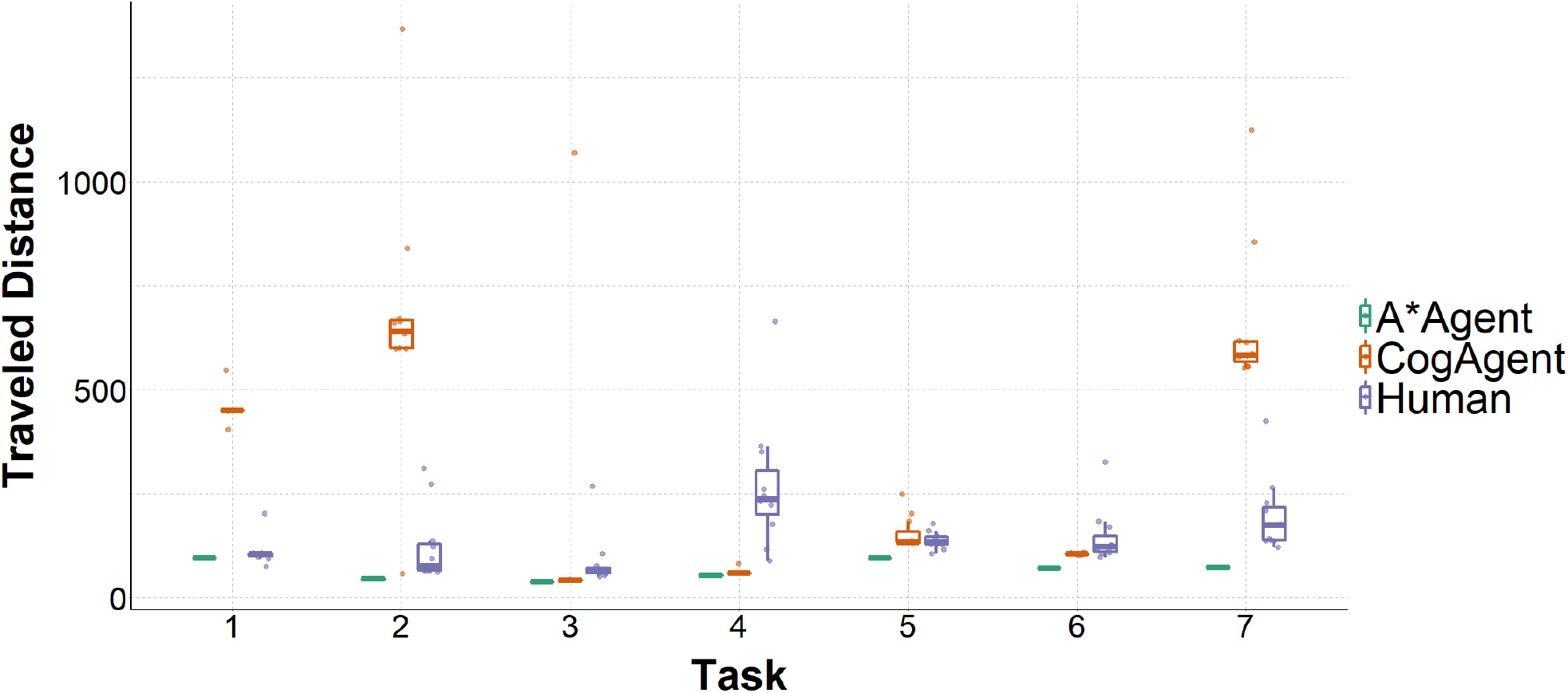
Average Distance Traveled by Human Participants, Uncertainty-based Cognitive Agents, and the A* Shortest-path Agent

In considering what features of the environment may have led to these discrepancies, we noted that the tasks in which the proposed uncertainty-based agent took relatively lengthy paths had fewer signs at crucial decision points (especially in tasks 2 and 7). This led to a greater number of wrong turns by both the humans and the uncertainty-based agents. However, the humans relied on other environmental cues and frequently backtracked in unpromising areas, whereas the agents, relying solely on the signage information for environmental cues, persistently continued to explore areas of the building that the humans rapidly realized were unfruitful. The “follow-your-nose” heuristic of the agents appeared to contribute to this behavior, prompting them to forge ahead in conditions of uncertainty. This heuristic did not consistently align with the human’s behavior of frequent backtracking. Task 4 was unique in that the humans took much longer routes on average than the uncertainty-based agent; however, this outcome appeared to also be a result of the “follow-your-nose” heuristic, combined with the “floor-first” strategy. In this task, agents moving ahead into unknown terrain quickly discovered a staircase that provided a short path to the destination, which was overlooked by most of the humans. We noticed a similarity in other multi-floor tasks, where the cognitive agents tended to find the correct floor very rapidly and humans spent greater time exploring incorrect floors (see supplemental material Figure S2). Overall, the distance travelled by humans tended to be closer to the shortest-path algorithm than to our uncertainty-based cognitive agent, but these differences varied greatly by task and appeared to be related to the broader route-seeking heuristics rather than to the impact of uncertainty levels.

## 5. Discussion and Conclusions

This research demonstrates the potential for improving indoor wayfinding simulation models by incorporating empirical evidence about human uncertainty levels in relation to environmental cues. Our Study 1 demonstrated that several signage-related indoor variables (most notably, the elapsed time from seeing a helpful sign) could be associated with changes in wayfinding uncertainty. However, the correlation of most of these variables with uncertainty was relatively weak, and more research is needed to evaluate how humans respond to complex environmental variables. Previous study suggests most intense uncertainty usually arises when the agent approaches a decision point, such as branching routes (Jonietz & Kiefer, 2017b). In addition to this route-choice uncertainty, we have incorporated on-route uncertainty into our existing framework. On-route uncertainty emerges when an individual is navigating between decision points. For example, when the person is walking down a long corridor or traversing a large open space and is attempting to confirm whether the current location is indeed along the correct route to their destination. This identified on-route uncertainty aligns with the concept of “confirmation bias” in psychology, suggesting that individuals tend to discount information that contradicts previous choices and seek confirmation to maintain confidence in their decisions. (Kappes et al., 2020; Klayman & Ha, 1987; Sharot & Sunstein, 2020). The simulation model that we constructed based on the findings of Study 1 was shown in Study 2 to be better at predicting human’s uncertainty responses during wayfinding compared to simple heuristic agents and random agents. In evaluating the distances travelled during wayfinding, however, we found a great deal of variability across different tasks between the human participants, our uncertainty-based agent, and a shortest-path agent. In some tasks the humans took a much longer path, and in other tasks the uncertainty-based agent took a much longer path. Our analysis of these outcomes suggested that the humans were responding to additional environmental cues beyond the ones evaluated in the current study, and that they were likely using more nuanced wayfinding heuristics. Thus, while the proposed uncertainty-based agent demonstrated more realistic behavior compared to a shortest-path algorithm, more research and computational development is needed to bring the agent closer into alignment with human navigational cognition.

### 4.1. Implications for Design

The goal of creating more realistic wayfinding models is to allow designers and planners to optimize human navigational efficiency and spatial experiences. One notable result from our Study 1 was that increasing navigational uncertainty was most closely correlated with the time since the wayfinder had last seen a helpful sign. This outcome indicates that designers should emphasize robust sign coverage throughout buildings, rather than only at notable intersections; a recommendation that aligns with some of the previous work in wayfinding design (Arthur & Passini, 1992). It is also important to note in this context that a “helpful sign” refers only to one that effectively communicates information to the user. Sign designers must therefore consider user demographics, including language backgrounds, as well as cognitive and physical abilities (Kalantari et al., 2022). In the specific area of wayfinding computation, our study is part of a larger movement toward integrating more robust data about human responses and behaviors in relation to environmental cues. The findings demonstrate how empirical measurements of uncertainty, combined with heuristic strategies, can be used to introduce more realistic behaviors compared to shortest-path algorithms. However, the findings also indicated that the agent’s performance was quite variable in relation to the humans’ performance on different wayfinding tasks, and that more robust research is needed to fine-tune these agent responses before they can become an effective diagnostic tool for designers.

### 4.2. Limitations and Future Work

While the current study offers a promising approach to modeling uncertainty in human wayfinding through the use of cognitive agents, it is crucial to acknowledge the work’s limitations and the need for further investigation. The current research employed a relatively small participant sample size, consisting of 39 participants (11 for the model creation data and 28 for the validation study). To ensure the generalizability of the findings, future work should aim to augment and diversify the sample, encompassing a broader range of participants and demographics. This will help to enhance the robustness of the model and its applicability to different populations. Additionally, the study focused exclusively on a single educational building as the environment for the wayfinding tasks, during both the model creation and validation stages. To augment the validity of the model, future research will eventually need to collect training data in more diverse indoor contexts and evaluate the outcomes in a similarly wide range of buildings. Using diverse settings such as hotels, transportation hubs, and healthcare facilities in addition to educational buildings will provide valuable insights into the generalizability of the model across different real-world contexts.

The current model evaluated a limited number of environmental factors, focused on visual signage, the number of available routes, and local isovist spatial variables as the potential sources of uncertainty (as well as an agent’s spatial abilities and prior knowledge of the environment). However, wayfinding is a multimodal process that may involve numerous other cues, such as the use of landmarks, auditory information, and the behavior of nearby individuals (Carlson et al., 2010). The omission of these broader variables is the likely reason why our agent’s wayfinding performance varied extensively in comparison to the human participants, and including a broader range of environmental cues may help to refine the agent’s performance.

An additional important undertaking for future research will be to improve the wayfinding heuristics that mediate the agents’ responses to environmental variables. While our simulation included a few basic heuristics (such as prioritizing searches for floor-level transition points), it does not fully reflect the role of cognitive processes, such as memory, attention, and problem-solving, in wayfinding behavior. Allowing the simulated agents to engage in more “reasoning” about the environment, while a daunting programming task, could serve to further enhance the model’s realism and utility (Wiener et al., 2009). One important direction in this regard is to examine and simulate the feedback loop between experiences of uncertainty and subsequent wayfinding behaviors, specifically focusing on information-seeking and backtracking. Our analysis suggested that the human participants did not consistently follow the “floor strategy” and “follow your nose” (continue straight ahead) heuristics, but instead engaged in more pausing, backtracking, and searching behaviors when confronted with uncertainty. By conducting more careful evaluations of these strategic responses, wayfinding simulations can better model the ways in which environmental cues and resulting states such as uncertainty translate into behaviors.

Finally, wayfinding simulations can benefit from developing a variety of agents with different abilities and proclivities, thereby helping designers to consider the experiences of people from different walks of life. The variables of spatial skills and prior environmental knowledge, as evidenced in the current research, can potentially be used to help diversify agents in this fashion. However, researchers should consider other human factors as well, including personality variables such as risk-taking and the presence of wayfinders who have perceptual differences such as visual impairments or language barriers. Considering all of these potential factors that can influence wayfinding experiences and outcomes, there is a tremendous range of opportunity for more evidence-based modeling work in wayfinding research.

## Notes

### Competing Interest Statement

The authors have declared no competing interest.

### Summary of Updates

In this revision of the manuscript, we have undertaken substantial updates to improve and validate our findings. Primarily, we have conducted a comprehensive data analysis involving all participants, thereby strengthening our results by including a more diverse dataset. In addition, we have incorporated a new validation study that applies our computational model in a real-world scenario. These updates help solidify our research claims and provide additional robustness to the conclusions drawn from our study.

