## Supplementary Material for "An Evidence-based Cognitive Model of Uncertainty during Indoor Multi-level Human Wayfinding"

**Figure S1.**

*Decision-making Zones of the Three Floors*

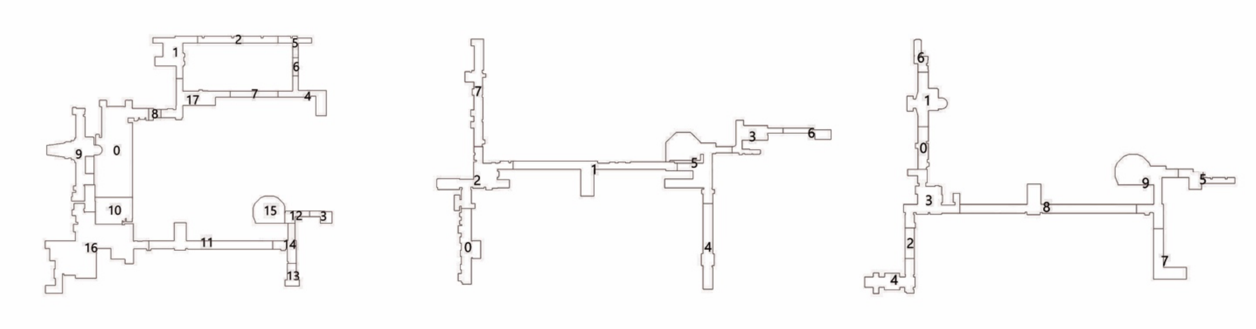


*Note: We divided the walkable areas of the building into distinct "decision zones". We defined key decision-making zones where participants must choose to stay on their current route, while the remaining areas, including long corridors, staircases, and dead ends, were classified as non-decision-making zones.*

**Table S1.**

*Full List of Isovist Metrics*

| | Isovist metric | Description |
|---|---|---|
| 1 | Area | Area of the isovist. |
| 2 | Distance Weighted Area | Area weighted by distance of each isovist ray. |
| 3 | Perimeter | Perimeter of the isovist. |
| 4 | Compactness | Ratio of the isovist area to the area of a circle with same perimeter as the isovist. |
| 5 | Circularity | Ratio of the square of the perimeter to area. |
| 6 | Convex Deficiency | Ratio of the area of the dent over the area of the hull. |
| 7 | Occlusivity | Length of occluding boundaries within the isovist. |
| 8 | Max Radial | Maximum length of radial lines. |
| 9 | Mean Radial | Mean length of radial lines. |
| 10 | Standard Deviation | Square root of variance. |
| 11 | Variance | Variance of the length of radial lines. |
| 12 | Skewness | Third moment about the mean of the radials. |
| 13 | Dispersion | Average radial distance divided by the maximum radial distance. |
| 14 | Elongation | Square root of the ratio of the eigenvalues of the covariance matrix from the isovist vertices. |
| 15 | Drift Magnitude | Distance between the observation point and the mass centre of an isovist polygon. |
| 16 | Drift Angle | Angle between the occupant’s facing direction and the mass centre of an isovist polygon. |

**Figure S2.**

*Uncertainty Heatmaps for Human Participants and Cognitive Agent by Task Number*

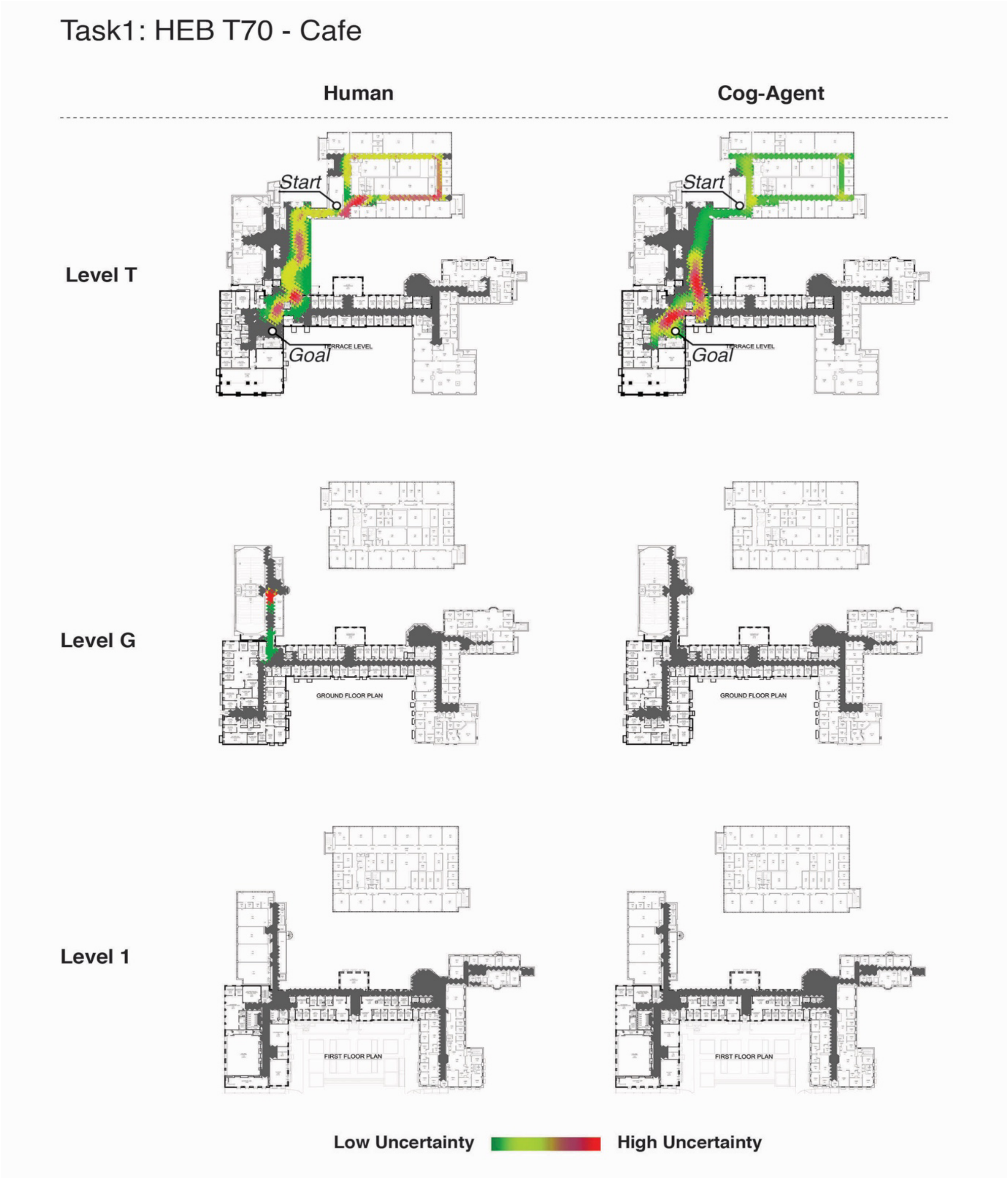


*Note: For this single-level task, the cognitive agent's spatial distribution of uncertainty experience resembles human participants in most of the areas.*

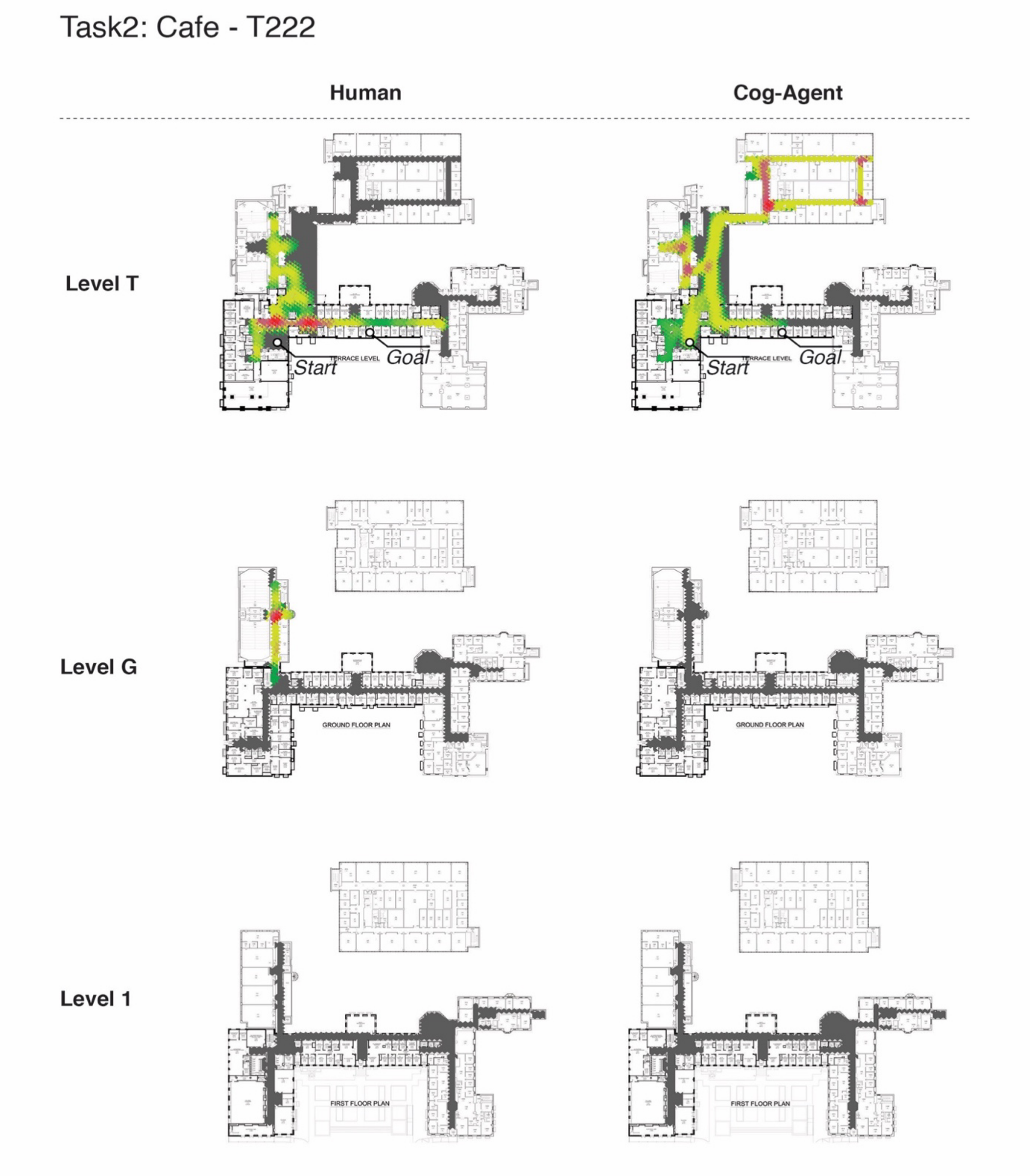


*Note: This is a short task with a low degree of sign coverage. The cognitive agents explored a much larger area than the humans because the agents didn't backtrack. Some humans missed the destination but then quickly reversed course.*

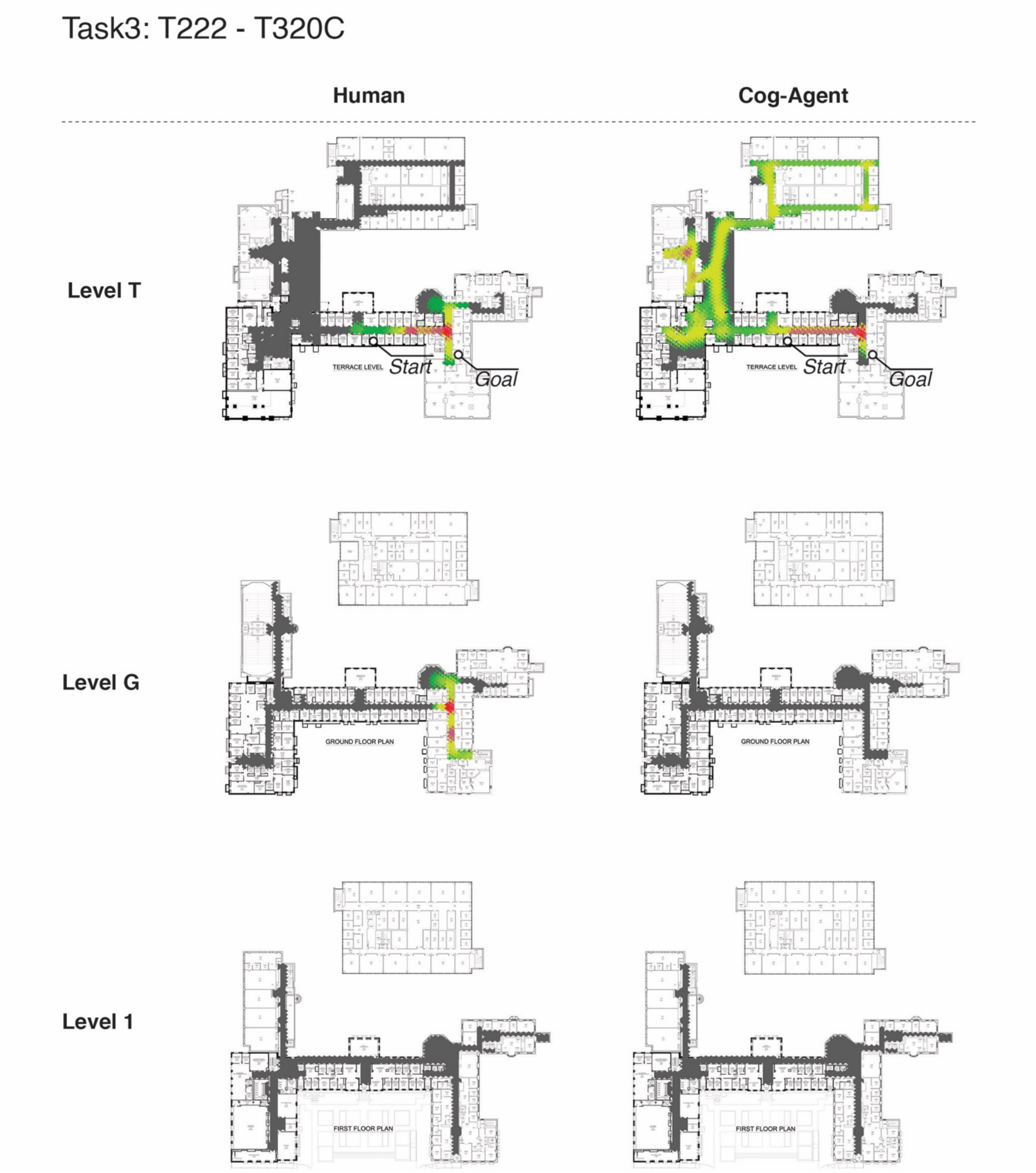


*Note: Most humans found the destination quickly but some went to the wrong floor. A few agents turned in the wrong direction at the beginning, but their uncertainty distribution generally resembles the humans.*

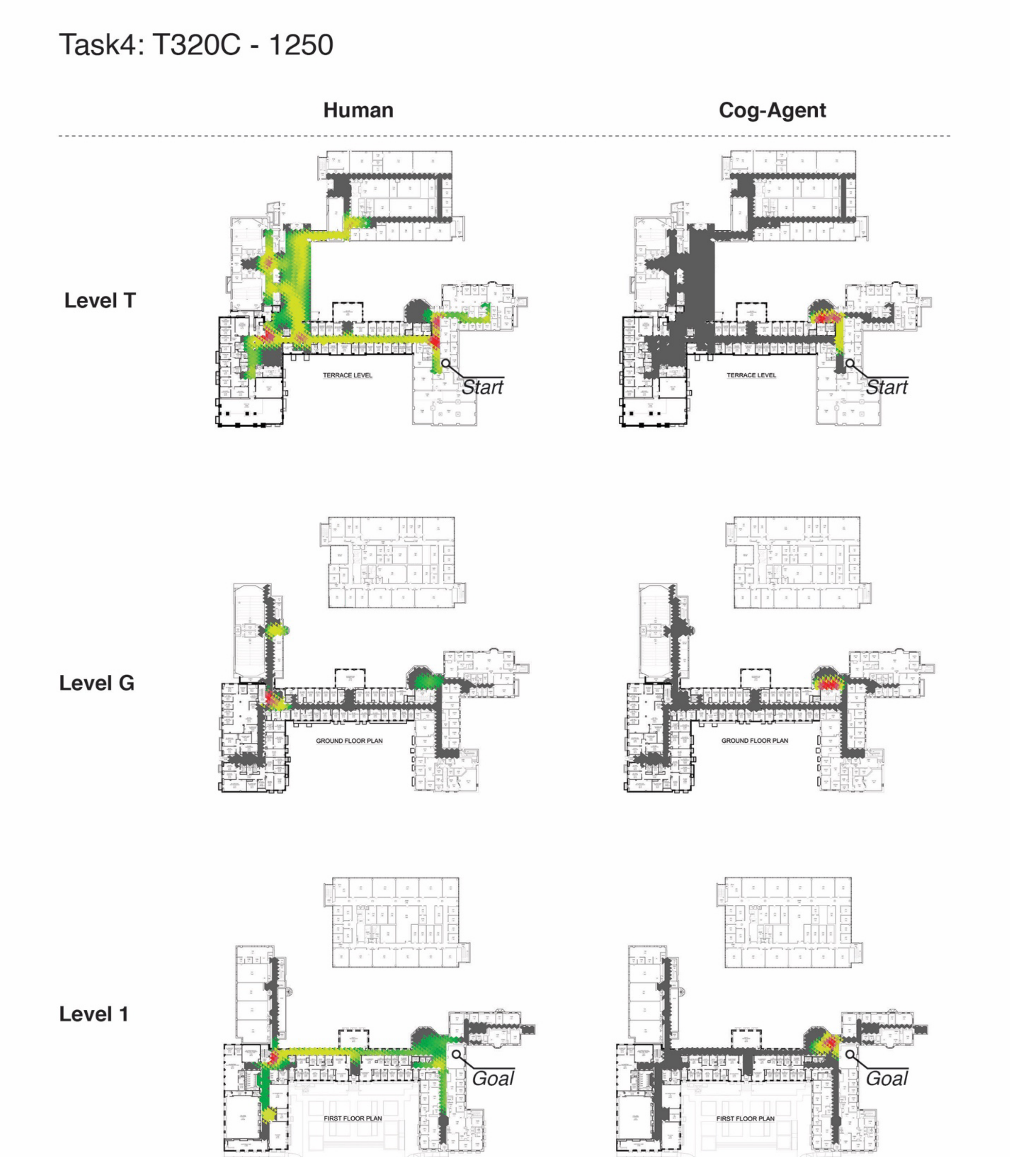


*Note: In this multi-level task, the agents found a shortest-path vertical staircase much sooner than the humans, presumably as a result of the agents' "follow-the-nose" and "floor strategy" heuristics. As a result the uncertainty distribution was very different from the human participants.*

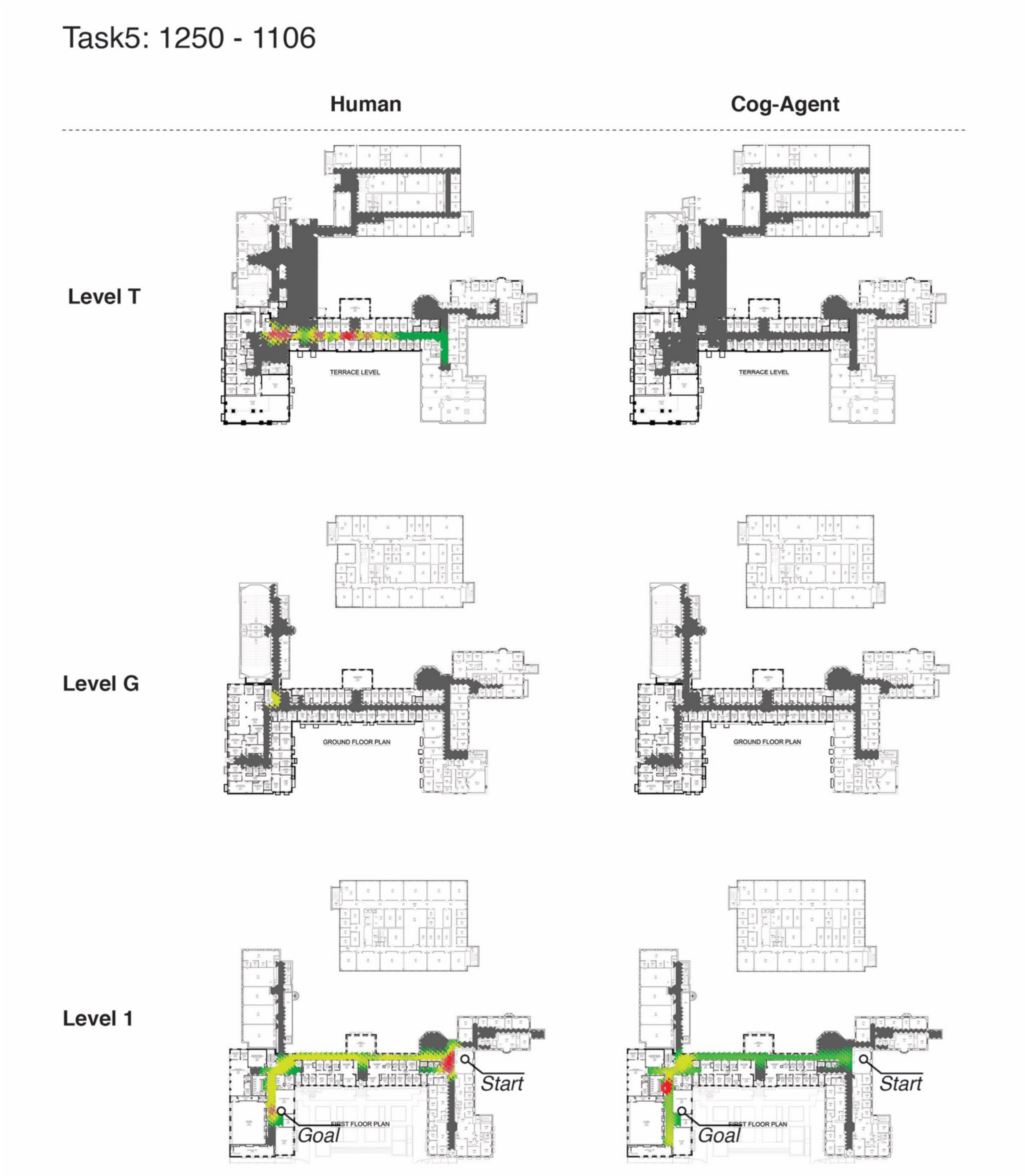


*Note: The agents' trajectory resembled the humans in this single-level task. However, human participants felt more uncertain at the beginning and at the end, presumably since the goal was blocked from view.*

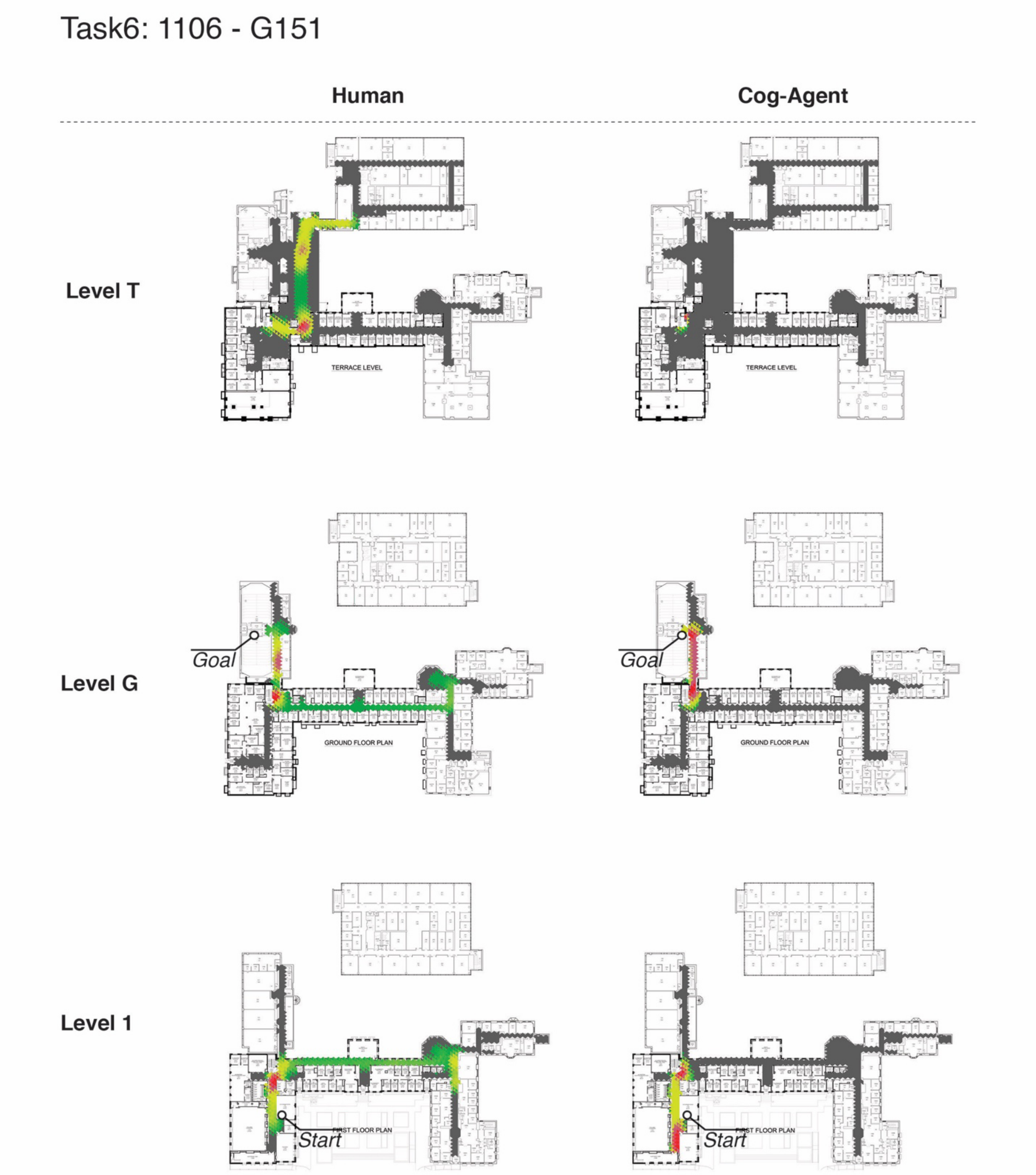


*Note: In this multi-level task, agents' uncertainty distribution and trajectories resemble the humans at level 1 and at level G. The agents' heuristics prevented them from taking a wrong path by entering Level T, while the humans spent significant time on that incorrect floor.*

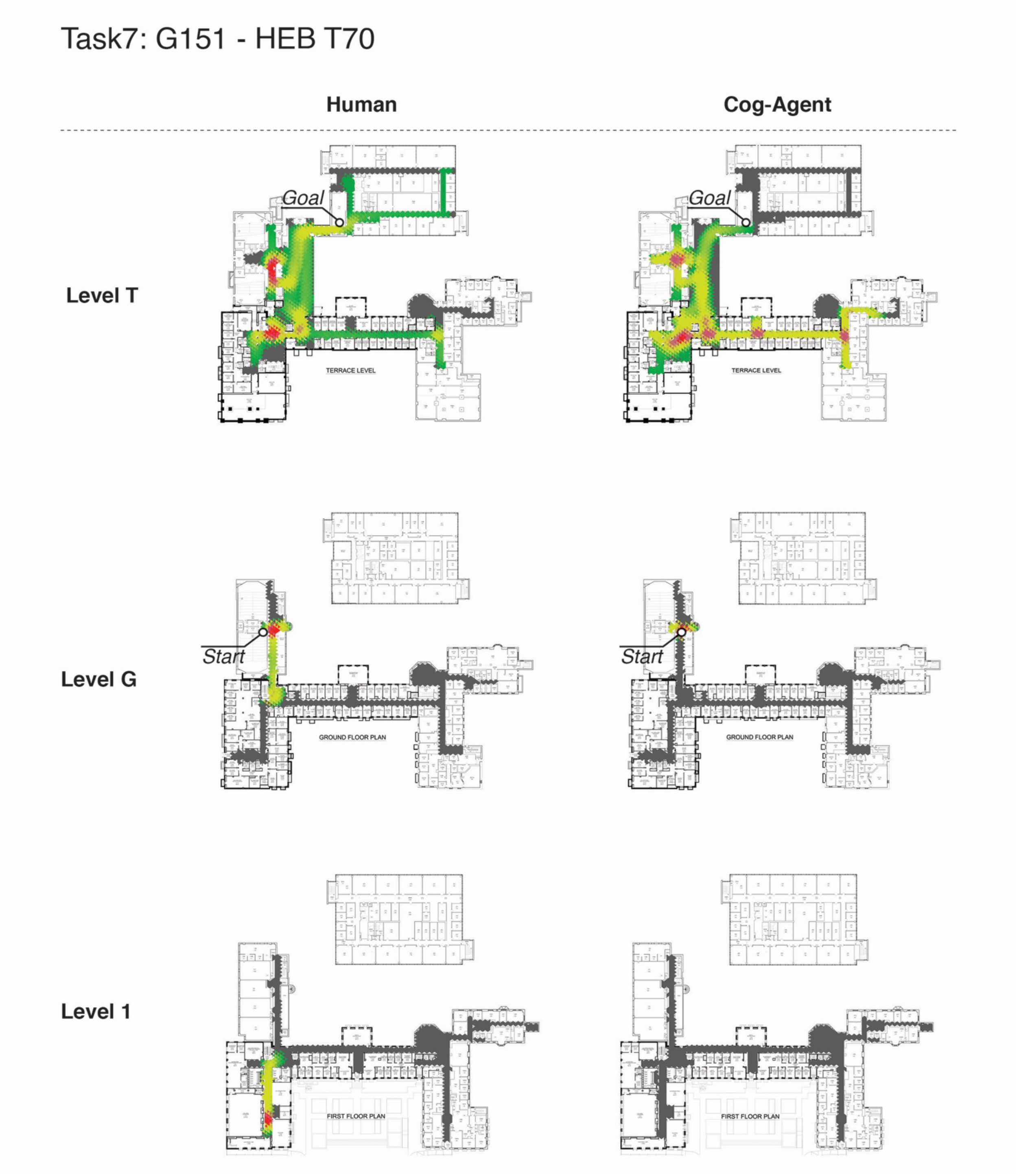


*Note: In this multi-level task, the agents' uncertainty distribution resembles the humans at level T. The humans spent more time exploring the other incorrect floors.*
